# Functional Genomics Screen with Pooled shRNA Library and Gene Expression Profiling with Extracts of *Azadirachta indica* Identify Potential Pathways for Therapeutic Targets in Head and Neck Squamous Cell Carcinoma

**DOI:** 10.1101/381749

**Authors:** Neeraja M Krishnan, Hiroto Katoh, Vinayak Palve, Manisha Pareek, Reiko Sato, Shumpei Ishikawa, Binay Panda

## Abstract

Tumor suppression by the extracts of *Azadirachta indica* (neem) works via anti-proliferation, cell cycle arrest, and apoptosis, demonstrated previously using cancer cell lines and live animal models. However, very little is known about the molecular targets and pathways that the neem extracts and the associated compounds act through. Here, we address this using a genome-wide functional pooled shRNA screen on head and neck squamous cell carcinoma cell line treated with crude neem leaf extracts, known for their anti-tumorigenic activity. By analyzing differences in global clonal sizes of the shRNA-infected cells cultured under no treatment and treatment with neem leaf extract conditions, assayed using next-generation sequencing, we found 225 genes affected the cancer cell growth in the shRNA-infected cells treated with neem extract. Pathway enrichment analyses of whole-genome gene expression data from cells temporally treated with neem extract revealed important roles played by the TGF-β pathway and *HSF-1*-related gene network. Our results indicate that neem extract simultaneously affects various important molecular signaling pathways in head and neck cancer cells, some of which may be therapeutic targets for this devastating tumor.

## Introduction

Despite previous knowledge on the use of the ethnobotanical database, evidence-based know-how and the mechanism(s) of action of many plant-derived compounds are currently lacking. *Azadirachta indica,* neem, is one such species that is unique, versatile and important. The neem tree is also one of the most intensively studied sources of natural products with plant-derived extracts showing anti-bacterial, anti-fungal, anti-viral, anti-inflammatory, anti-hyperglycaemic, immunomodulatory, anti-malarial, and anti-carcinogenic properties (reviewed in (Subapriya and Nagini, 2005). Past efforts on research on neem have mostly been focused on pre-clinical studies, concentrating mainly on two purified neem-derived compounds, nimbolide and azadirachtin, due to their pharmaceutically- and agriculturally-important properties, respectively. The emergence of genome-wide tools and techniques has opened up new avenues to pursue research through systematic studies on cancer drug target discovery using neem-derived metabolites. One such method is RNA interference. It is a powerful and proven system to perturb gene function in higher organisms by phenotypic screening, with potential ability to identify pathways and networks of genes involved in cancer (Tang et al., 2008; Blakely et al., 2011; Poell et al., 2011; Iorns et al., 2012; Mendes-Pereira et al., 2012; Poell et al., 2012; Singleton et al., 2013). Use of pooled genome-wide libraries of short hairpin RNA (shRNA) coupled with next-generation sequencing offers higher sensitivity and broader dynamic range to screen for biological activity and therefore has the potential to identify the pathways for drug actions through network-based approaches (Erler and Linding, 2010; 2012).

Head and neck squamous cell carcinoma (HNSCC) is the sixth leading cause of cancer worldwide (Jemal et al., 2011) with a 5-year survival of less than 50% (Mishra and Meherotra, 2014). In the past, the ability of neem-derived compounds to stop cell proliferation and their mechanism(s) of action have been reported, including in HNSCC cell lines (Roy et al., 2007; Harish Kumar et al., 2009; Harish Kumar et al., 2010; Priyadarsini et al., 2010; Gupta et al., 2013; Hsieh et al., 2015; Kasala et al., 2015; Patel et al., 2016; Chien et al., 2017; Kowshik et al., 2017). However, efforts towards finding novel drug targets and the pathways using genome-wide tools for neem-derived compounds is currently lacking. The current study combined genome-wide shRNA based functional screening and whole-genome gene expression assay using a head and neck cancer cell line, paving ways to characterize and understand the critical molecular pathways for the therapeutic properties of the neem leaf extract against HNSCC.

## Materials and Methods

### Neem plant extracts

We tested leaf, bark, fruit, dry seed and twigs from neem plants to make ethanolic extracts. First, the plant parts were collected fresh and thoroughly washed with MilliQ water followed by 70% ethanol and the process was repeated thrice with a final wash with MilliQ water before they were air dried under controlled condition for several days. Once the plant organs were dry, 250gm of each organ was measured and ground separately using a grinder, mixed with 500ml of ethanol and transferred to a glass bottle. The mixture in the glass bottle was rocked overnight to mix thoroughly and then centrifuged at 5000g for 10min. The supernatant was separated, and the residue at the bottom was again treated with 500ml of ethanol with overnight rocking at room temperature followed by centrifugation. The process was repeated three times, and the supernatant was pooled from all the three spins before proceeding to the next step. The ethanolic extract (referred as neem extract) was evaporated using an oven at 55^0^C till the materials became very viscous and gooey like and then kept at 4^0^C until further use.

### Reconstitution of neem extracts

Neem extracts were dissolved in DMSO to make a stock solution, which was serially diluted further with Dulbecco’s Modified Eagles’ Media (DMEM) to make a final concentration of 2mg/ml as working stock solution. The working stock solution was passed through a 0.22micron sterile filter and stored until further use. The filtered and sterile stock solution was further diluted with DMEM to make the final solution and was used for treatment.

### Cell culture and IC_50_ calculation

Multiple human HNSCC cell lines (UPCI:SCC029B and UPCI:SCC040, both gifts from Dr.Susan Gollin, University of Pittsburgh, PA, USA (White et al., 2007); UM-SCC47, a gift from Dr. Thomas Carey, University of Michigan, MI, USA (Brenner et al., 2010); HSC-3 and HSC-4, purchased from RIKEN, Japan (Momose et al., 1989) were tested with all neem extracts before HSC-4 was chosen for further study as the cells showed excellent growth inhibitory pattern in the presence of neem extracts. All the cells were maintained in DMEM supplemented with 10% FBS, 1X MEM non-essential amino acids solution and 1X penicillin/streptomycin mixture (Gibco) at 37^0^C with 5% CO_2_ incubation.

For IC_50_ calculation, the xCELLigence Real-Time Cell Analysis (RTCA) DP instrument (Acea Bio, USA) was used. The device monitors and quantifies cell proliferation, morphology change, and provides an opportunity to observe the quality of cell attachment real-time. HSC-4 cells (1.0 x 10^4^) were seeded onto wells with varying concentrations of neem extract (25-800μg/ml) for varying times (0-48hrs). The cells were kept under observation, and IC50 values were calculated for all extracts and those with DMSO only used as vehicle control. Neem leaf extract showed the best dose response (Fig. S1) and therefore chosen for further study. All subsequent experiments were carried out with the IC_50_ concentration of neem extract unless otherwise specified.

### Whole genome gene expression assays and data analyses

HSC-4 cells were seeded and treated with neem leaf extract (200μg/ml) at nine pre-defined time points (5min, 10min, 15min, 30min, 1hr, 3hr, 6hr, 10hr, and 24hr) and rescued with fresh complete medium post 48-hour treatment at these time-points. The samples were assayed for gene expression using Illumina whole-genome HumanHT-12 v4 expression BeadChip (Illumina, San Diego, CA), following the manufacturer’s specifications. RNA quality was checked using Agilent Bioanalyzer with RNA Nano6000 chip for integrity before being used in the gene expression assays. The RNA samples were labeled using Illumina TotalPrep RNA Amplification kit (Ambion, USA) and processed following the manufacturer’s recommendations. Gene expression data was collected using Illumina’s HiScan and analyzed with the GenomeStudio (v2011.1 Gene Expression module 1.9.0), and all assay controls were checked to ensure the quality of the assay and chip scanning. Raw signal intensities were exported from GenomeStudio for pre-processing and analyzed using R further. Gene-wise expression intensities from GenomeStudio were transformed, normalized using VST (Variance Stabilizing Transformation) and LOESS methods, respectively, using the R package lumi (Du et al., 2008) and further batch-corrected using ComBat (Johnson et al., 2007). The preprocessed intensities were subjected to differential expression analyses using the R package, limma (Ritchie et al., 2015) and fold change values were computed for various time-points in the treatment and rescue conditions versus the control samples. Genes that were serially up-regulated or serially down-regulated across at least four consecutive time-points were short-listed for further analysis.

### Genome-wide shRNA lentivirus library screening and transduction

We used a genome-wide shRNA library for signaling pathway genes consisting of 27,500 shRNAs (Human Module 1, Cat No DHPAC-M1-P from Cellecta) for our experiment (Figure 1). Plating of the cells, transfection, DNAse treatment, viral titer estimation, lentiviral transduction with polybrene and final puromycin selection were carried out following the manufacturer’s instructions. HSC-4 cells were infected with the shRNA lentivirus library and cultured with or without neem extract treatment and harvested, and frozen, followed by the extraction of their genomic DNA (Figure 1). Two concentrations of neem leaf extract (200 μg/ml and 300 μg/ml) were used for the treatment of cells. The RFP signals at the time of harvesting (72 hr after the lentivirus infection and before the puromycin treatment) were confirmed by observing the cells under a fluorescent microscope (Figure S2).

**Figure 1.**
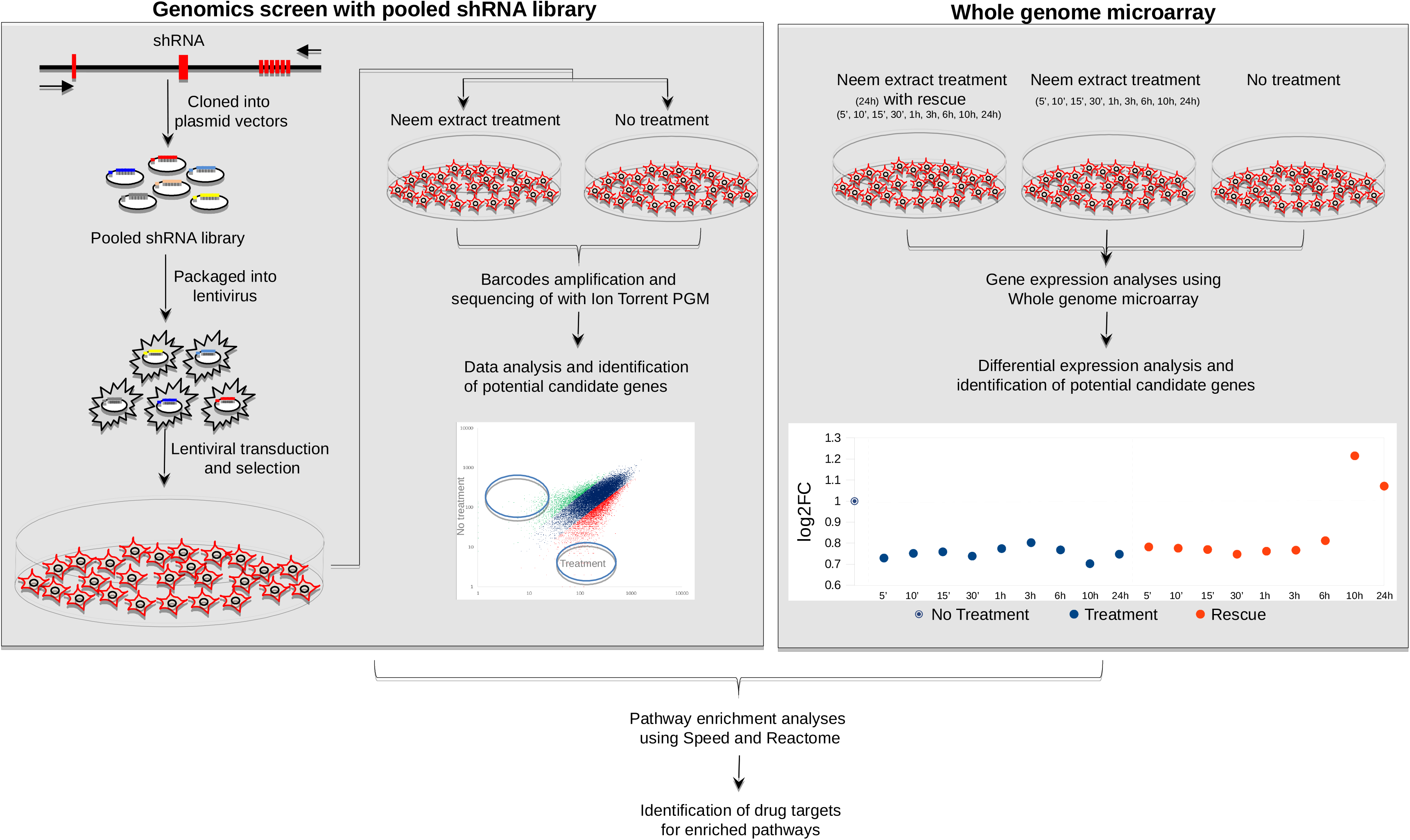
Overview of the screening method. A. Whole-genome functional shRNA screening experiment, B. Whole-genome gene expression microarray experiment.

### Next-generation sequencing post-genome-wide shRNA screening

We performed PCR amplification of barcode regions of the lentivirus library, followed by sequencing using the PGM system (Ion Torrent, Thermo Fisher, USA). Read QC was performed to meet the quality criteria for the reads in both control and shRNA library treated samples. The shRNAs with increased (> 2-fold) clone size in neem extract-treated cells compared to the control cells (with DMSO vehicle control) implied that the corresponding shRNAs contributed to the favorable cell survival or growth when treated with neem extract (Figure 2). On the other hand, the shRNAs whose clone sizes were decreased (< 2-fold) in neem extract-treated cells compared to the control cells implied that the knockdown of the corresponding genes worked in synergy with neem leaf extract for unfavorable cell survival or growth (Figure 2). Genes represented by at least two shRNAs were considered for further analyses to secure the reproducibility of the phenotypes of shRNA knockdown.

**Figure 2.**
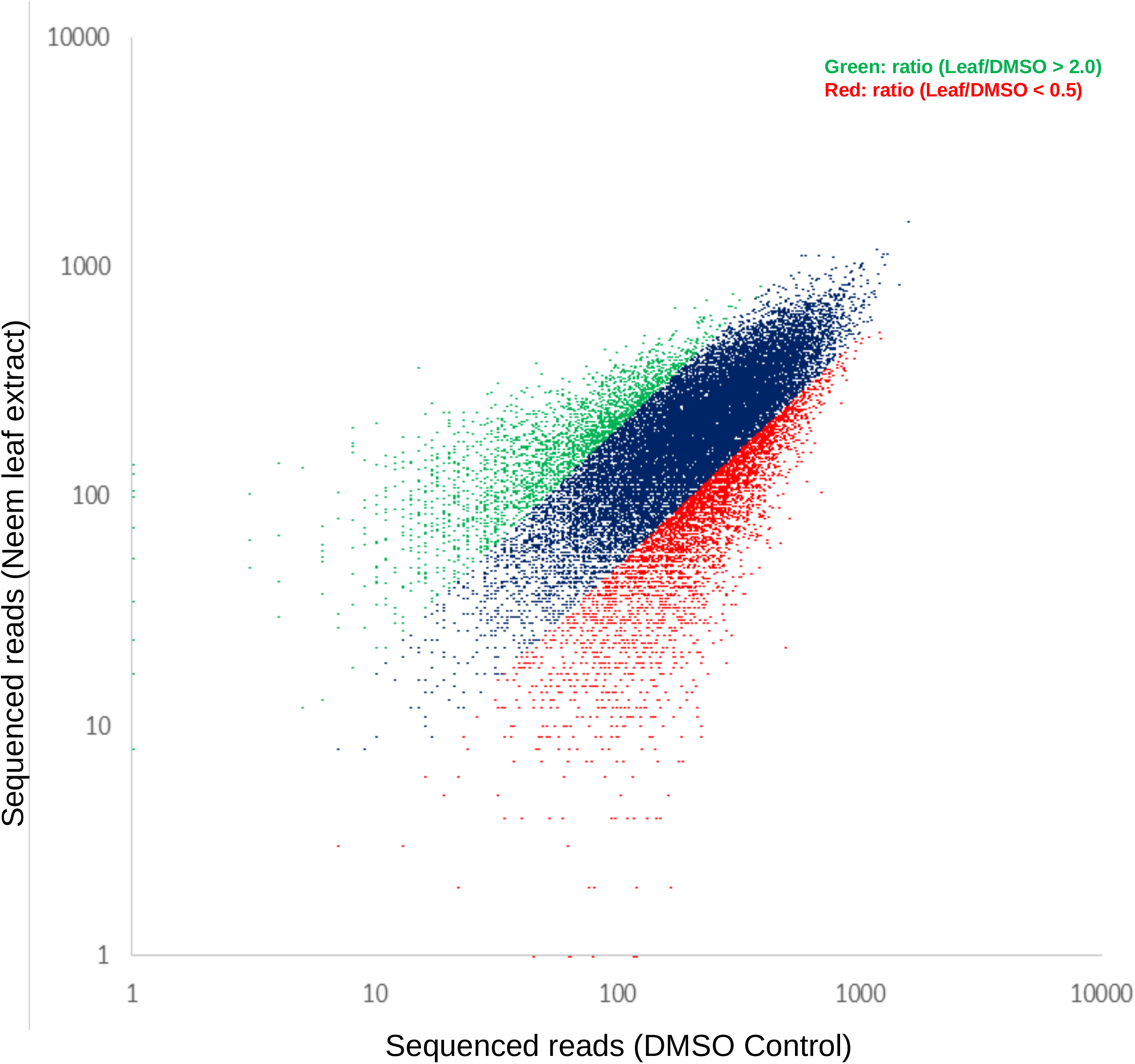
Scatter plot for barcodes in the sequencing reads in treated vs control neem leaf extract.

### Interaction network, drug targets and pathway enrichment analyses

The short-listed genes from the whole genome expression assay and the consensus list of genes derived from the neem-treated shRNA transfected cells were combined after removal of redundancy if any. These genes were subjected to pathway enrichment analyses using two independent resources: namely, Speed (Parikh et al., 2010) and Reactome (Croft et al., 2014). Speed enables discovery of upstream over-represented signaling pathways responsible for observed differential expression changes in the genes, and Reactome allows merging of pathway identification and over-representation while allowing interactors to increase the analysis background. We ran Speed while exercising the ‘Signature genes must be unique’ option to identify the significantly enriched pathway(s) first, before running it without choosing this option. This provided us with a complete list of genes perturbed in the enriched pathway(s). The perturbed list of genes from Speed and the over-represented genes and their interactors from enriched pathways detected by Reactome were subjected to network analyses using cBioPortal (Gao et al., 2013) (www.cbioportal.org) along with 50 significant most affected genomic profiles (mutations, putative copy number alterations from GISTIC and differentially expressed mRNAs (more than 2-fold) in the TCGA HNSCC dataset (provisional dataset)). Reactome was run by exercising the ‘Include Interactors’ option to increase the background list of genes. Cancer drugs were mapped to this network while differentiating between the FDA approved ones from those in the pipeline.

## Results

### IC_50_ calculation of the neem extracts on HNSCC cell line HSC-4

The IC_50_ values for the leaf, bark, fruit, seed and twig extracts were 0.184, 0.226, 2.932, 0.530, and 0.283 mg/ml, respectively, and the leaf extract showed the best dose response (Figure S1). Therefore, the leaf extract was used in subsequent experiments at IC_50_ concentrations unless otherwise specified.

### Gene expression changes in response to neem extract

The overall approach using a combination of functional genomics screening assay with the genome-wide shRNA library and the whole genome expression assay with treatment and rescue conditions is depicted in Figure 1. To precisely clarify the mode of molecular actions of the neem extract in a time-course manner, we investigated the changes in mRNA expression levels of all genes during the neem treatment compared to the steady state (24hr time-course), as well as their behaviors during the following rescue periods (24hr time-course).

Expression values for all genes at various time points (5min, 10min, 15min, 30min, 1hr, 3hr, 6hr, 10hr, and 24hr) and the follow-up rescue experiment (for the same periods of time) along with the fold change statistics and pathways affected is provided in Table S1. Expression of 28 genes was altered compared to no treatment, and their expression did not change significantly from the last time-point (24 hr) of neem leaf treatment (standard deviation across the last treatment time-point and in all rescue time-points <= 0.01). Out of those, 8 genes were temporally up-regulated *(CYP46A1, PDGFD, HOXA3, GRIK1, TRIM10, DDA1, OAZ1* and *NAT8B)* (R >= 0.6), and 20 temporally down-regulated *(KLK12, PQLC1, LEF1, IL15RA, CHAC2, LMNA, EEF1A1*, *EHD4*, *CS, COG5, MS4A6A, FH, PRPS1, GR14, HNRNPM, OPRM1, KCNIP4, KCNC4, IMPACT* and *MRAP*) (R <= −0.6). We selected genes (*n* = 98, 93 up-regulated *and* 5 down-regulated) with altered expression (ratio at least below 0.8 or at least above 1.2) over at least 4 consecutive time-points (Table S2) for pathway analysis comprising 15 cancer-related pathways (Table S1) for both treatment and rescue conditions (Figure 3). Genes involved in JAK-STAT, mTOR, VEGF, NF-κβ, and WNT signaling pathways, displayed a cumulative up-regulation effect over time, the action of neem extract was reversed upon the rescue. Their maximum values for expression change (log_2_fc) were 0.75, 0.9, 0.6, 0.75 and 0.4, respectively. Genes involved in the ECM-receptor interaction, on the other hand, were cumulatively downregulated, that was completely reversed upon the rescue (Figure 3).

**Figure 3.**
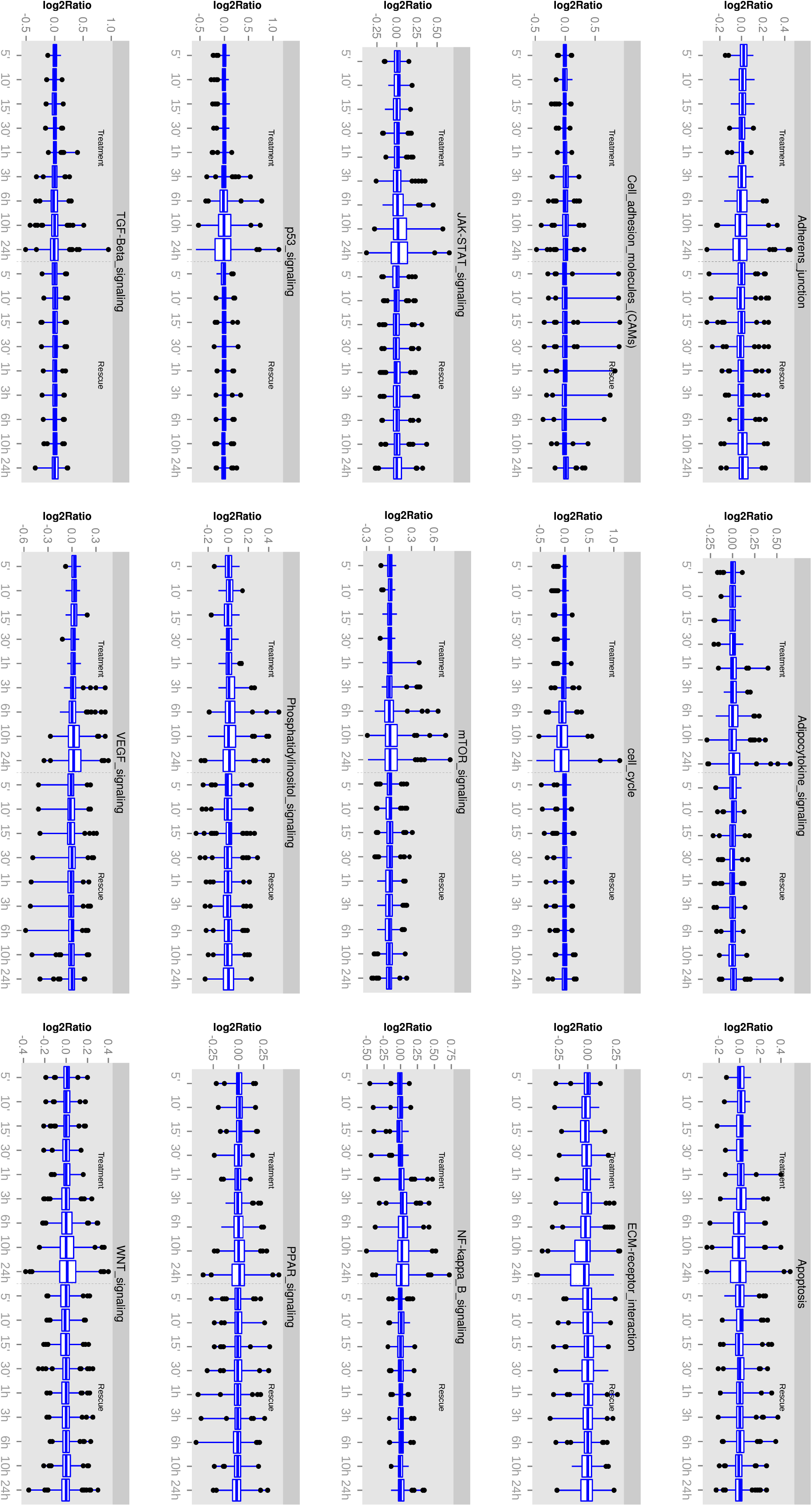
Cancer-related genes and pathways affected by neem extract at various time points and with the follow up rescue.

### Functional genomics with pooled shRNA library with neem extract treatment

A representative result of the shRNA screening is shown in Figure 2. Sequencing read statistics, and unique shRNA reads obtained after next-generation sequencing post neem extract treatment are provided in Table 1. The results from the PCR amplification of the barcodes, in control and treated cells, showed 1369 and 1011 shRNAs with high (ratio versus control: >2) and with low (ratio versus control: <0.5) clonal sizes respectively (Figure 2). This represented 149 and 78 genes, respectively, where more than one shRNA were involved (Table S4). The direction of changes in clonal size for genes are described in Table S4. To globally obtain a set of genes whose function and/or expression were somehow related to the neem extract treatment, we investigated the gene lists from the two independent screening experiments.

**Table 1.** Sequencing read statistics in functional pooled shRNA screening experiments.

|  | Total number of reads | Useful reads | % | Number of shRNA unique hits (total 27,495 shRNA) | % |
| --- | --- | --- | --- | --- | --- |
| DMSO control for neem leaf extract | 7,658,211 | 6,762,717 | 88.3 | 27,260 | 99.1 |
| Neem leaf extract (200µg/ml) | 6,727,689 | 6,000,047 | 89.2 | 27,344 | 99.5 |
| Neem leaf extract (300µg/ml) | 7,620,025 | 6,621,474 | 86.9 | 27,397 | 99.6 |

### Interaction network of genes and pathways picked from the gene expression and shRNA screening assays

We combined the short-listed genes whose expression were significantly changed by the neem extract over at least 4 consecutive time-points (Table S2) and those derived from the functional shRNA screening, resulting in a consolidated list of 321 non-redundant genes (Table S5). These genes were considered to be affected somehow by the neem extract treatment either or both in the functional and/or expression aspects; thus, we further used this list for pathway and interaction network analyses to identify any molecular pathways influenced by the neem extract.

The list of 321 genes (Table S5) compiled from the combination of the serially up- or down-regulated genes across at least four consecutive time-points (3hr, 6hr, 10hr and 24hrs) with large expression change (ratio at least below 0.8 or at least above 1.2) and genes with significant changes in the clonal sizes of shRNA-infected cells in the neem extract treatment, was subjected to pathway enrichment analyses, using Speed and Reactome analysis tools independently. The enrichment analyses using Speed while investigating only the unique set of signature genes resulted in only one pathway, the TGF-β signaling pathway *(p* = 7.67e-06; FDR = 2.87E-05), with a significant FDR adjusted P value at the 95% level of confidence (Figure 4A). The unique genes perturbed in this pathway were *TGIF1, SPHK1, DDIT3, RGS16, LRRC15, VDR* and *GABARAPL1.* When all genes in the pathways were investigated, all signalling pathways passed the FDR significance threshold, with MAPK_PI3K signalling pathway at the top, with the least FDR at 3.53e-13 and 18 genes perturbed out of 118 background genes, closely followed by TGF-β at an FDR of 4.69e-13 and 19 genes perturbed out of 142 background genes (Figure 4B). The pathway enrichment analyses using Reactome resulted in three pathways with significant FDR (< 0.05): Attentuation phase (FDR = 3.56e-05), HSF-1-dependent transactivation (FDR = 4.4e-05) and *HSF-1* activation (FDR = 0.013) (Table S6). The genes over-represented in the Attenuation Phase pathway network were a subset of the two other *HSF-1* -related pathways; therefore, the *HSF-1* -related pathway was considered to be the enriched pathway in relation to the neem extract treatment. The perturbed genes common to both HSF-1-related pathways were *DNAJB1, HSPA1L, HSPA1A,* and *DEDD2. HSPB8* was unique to the HSF-1-dependent transactivation pathway network, while *RPA1* and *RPA2* were unique to the *HSF-1* activation pathway. *BAG3, PLK1,* and *SPHK1* were the interactors used in the analysis in the case of *HSF-1* activation pathway (Table S6).

**Figure 4.**
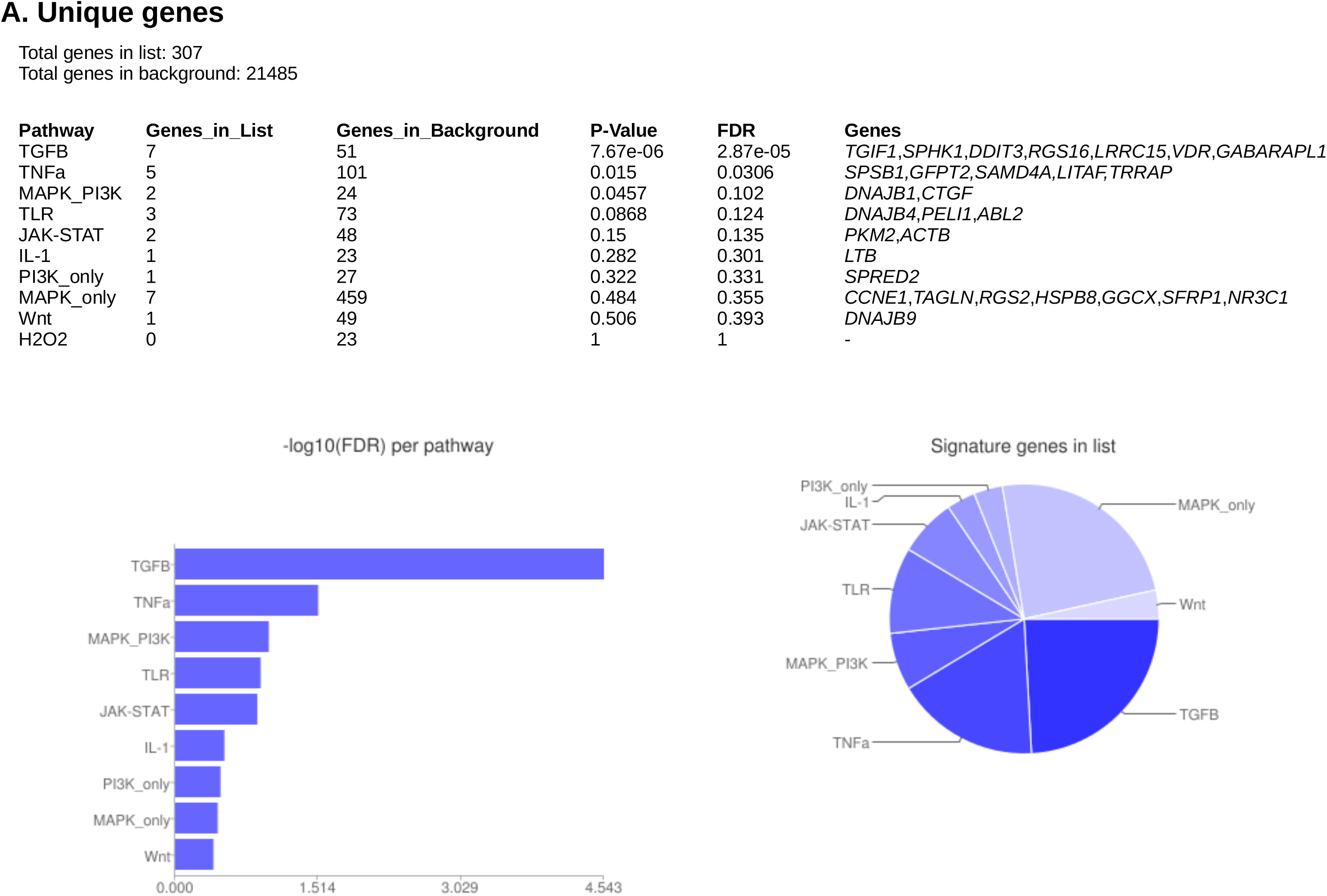
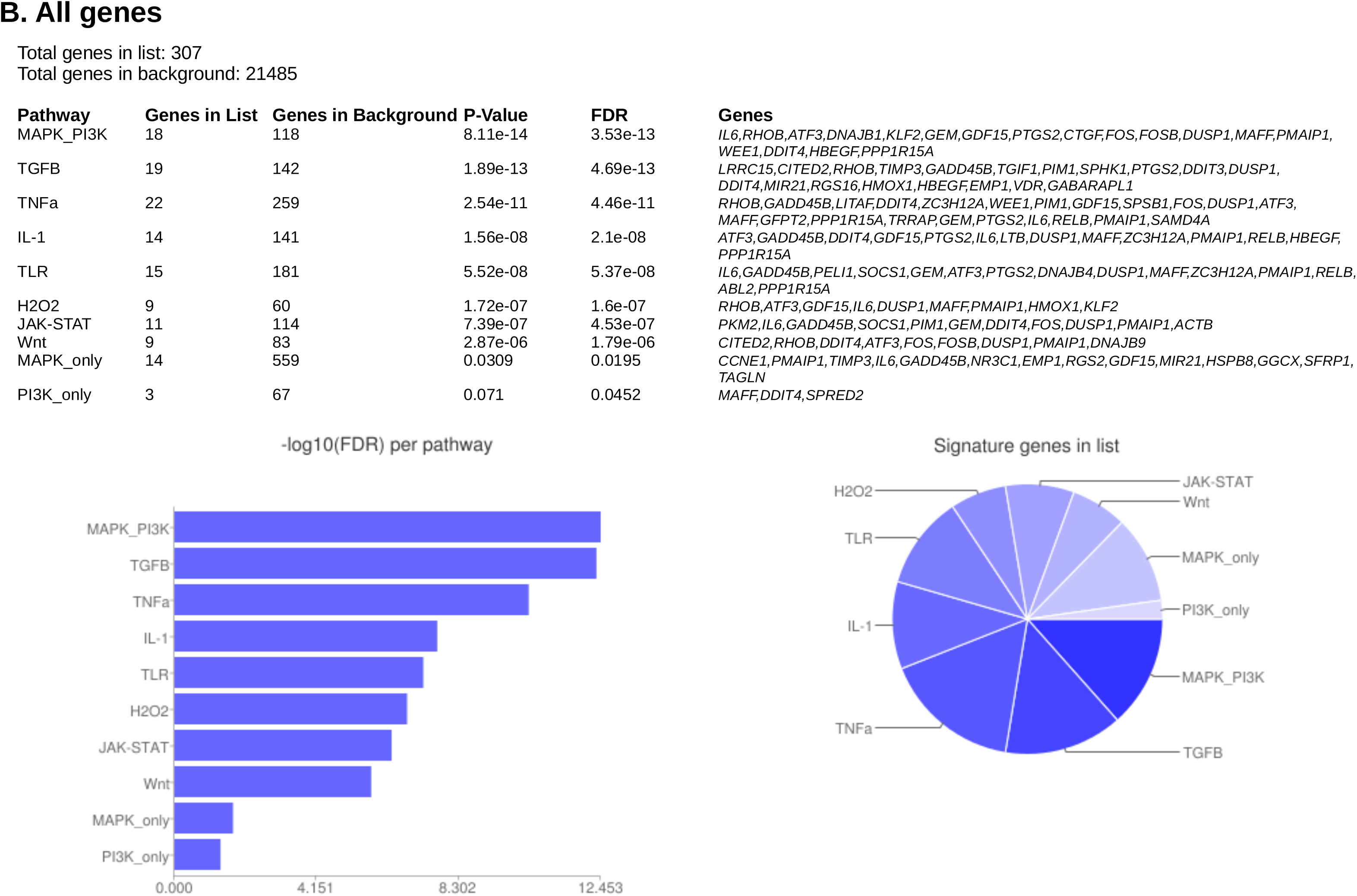
Enriched pathways and over-represented genes analyzed by Speed using the options: A. Unique genes and B. All genes.

### Identifying potential drug targets for neem extract

Finally, we explored any candidate molecular targets that may work in synergy with the neem extract for HNSCC patients. Depending on the list of genes in the pathways which were considered to be related to the mode of actions of neem extract (Figure 4), we obtained the network and drug target interaction information for the 19 perturbed genes *(LRRC15, CITED2, RHOB, TIMP3, GADD45B, TGIF1, PIM1, SPHK1, PTGS2, DDIT3, DUSP1, DDIT4*, *MIR21, RGS16, HMOX1, HBEGF, EMP1, VDR* and *GABARAPL1)* from the TGF-β signalling pathway as detected by Speed (Parikh et al., 2010), the 10 genes *(DNAJB1, HSPA1L, HSPA1A, DEDD2*, *HSPB8, RPA1, RPA2, BAG3, PLK1* and *SPHK1)* from the HSF-1-based pathways as detected by Reactome, (Croft et al., 2014) and the 50 most altered genes for the TCGA HNSCC provisional data from the cBioPortal (Gao et al., 2013) (www.cbioportal.org). A network was drawn using cBioPortal contained 55 nodes, including 16 out of 28 query genes (Figure 5). Thirty-one percent of the network was categorized as ‘drug-target interactions’. Among these networks, *PTGS2, VDR, RHOB, PIM1, HSPA1L, HSPA1A,* and *SPHK1* were used as targets for cancer drug development, previously. Out of these, *RHOB, VDR,* and *PTGS2* were the only candidates with FDA-approved anti-cancer drugs, with one each for *RHOB* and *VDR,* and 20 for *PTGS2* (Table S7). For others, quercetin (targeting *PIM1)* underwent seven clinical trials; and apricoxib (targeting *PTGS2)* underwent five clinical trials. Other genes such as *HMOX1* were the targets of 8 non-anti-cancer non-FDA-approved drugs (Table S7).

**Figure 5.**
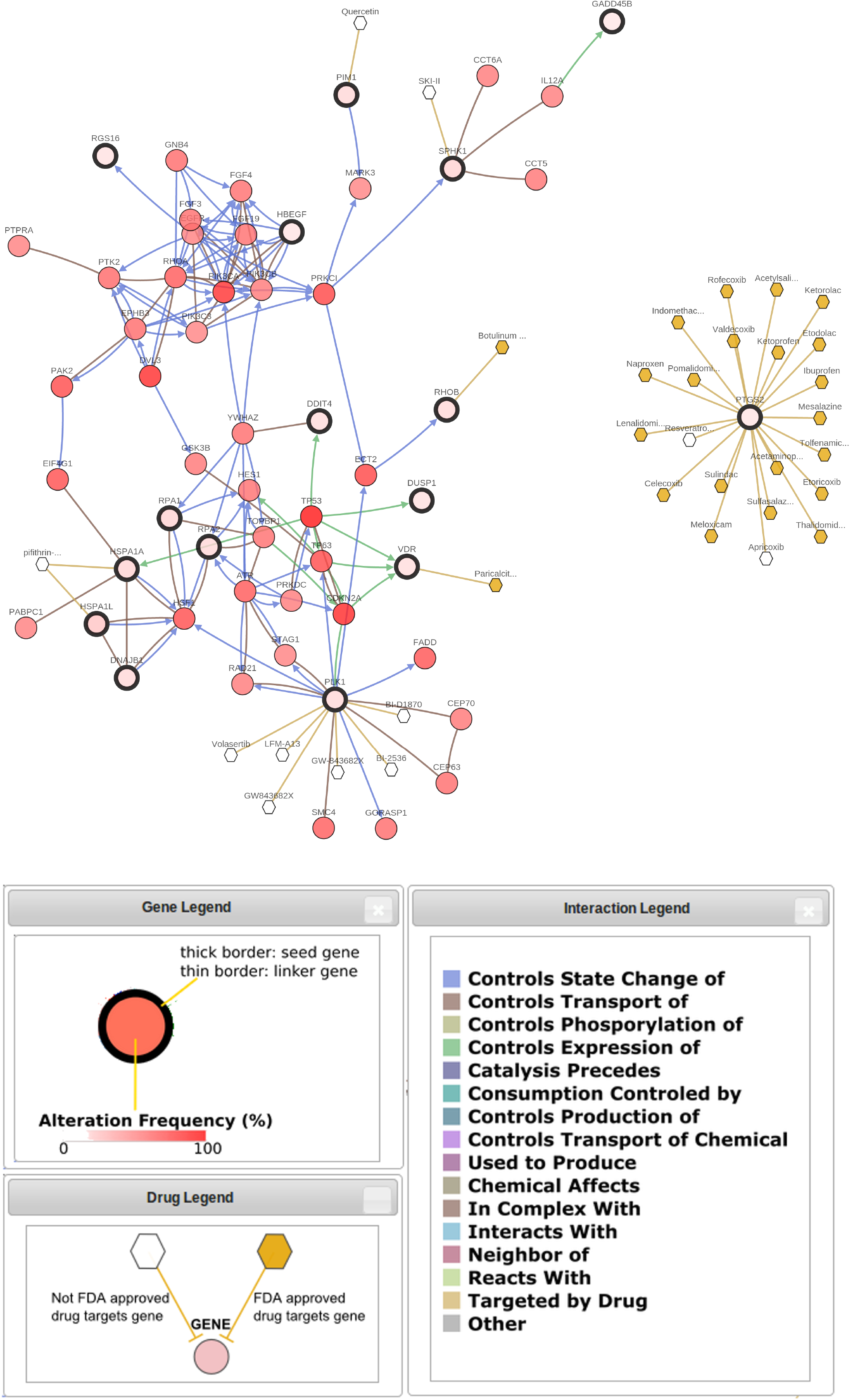
Hypothetical gene-drug interaction network in relation to the effect of neem extract in cancer.

## Discussion

Natural products are an important source of anti-cancer compounds (Hollosy and Keri, 2004). Neem leaf extract and derived metabolites are shown to act as anti-cancer compounds specifically through the modulation of expression of critical cancer-related signalling molecules, including in oral cancer animal models (Harish Kumar et al., 2009; Paul et al., 2011; Hsieh et al., 2015; Kasala et al., 2015; Patel et al., 2016; Kowshik et al., 2017). Additionally, it is shown to act through modulation of the proinflammatory microenvironment in colorectal cancer (Gupta et al., 2013). Purified neem compounds have been shown to induce cytotoxicity and apoptosis, and affect cell cycle in various cancer cell lines (Roy et al., 2007; Priyadarsini et al., 2010; Chien et al., 2017), decrease tumor incidence and pre-neoplastic lesions in live hamster models of oral oncogenesis (Harish Kumar et al., 2010). This study aimed to identify the molecular function of the neem extract with a particular focus on its anti-HNSCC property.

A critical application of the RNA interference-based functional screening is the identification of drug mechanisms as well as the synergic drug target(s); on the other hand, global time-course gene expression profiling under drug treatment also provides us significant insights into the drug mechanisms. In the current study, we combined these two independent global experiments to investigate the molecular mechanisms of the neem extract with focuses on the anti-cancer properties for HNSCC.

Pathway enrichment analyses were conducted for shortlisted genes which were considered to be differentially affected somehow by the neem extract, either or both in functional and/or gene expression aspects, identifying a possible role for the TGF-β signaling pathway and the *HSF-1* activation network. The *HSF-1* is related to the TGF-β signaling pathway and connects the IGF, TGF-β and cGMP pathways, and controls development processes (Barna et al., 2012). The precise molecular interactions and biochemical downstream effects of the neem extract in combination with these pathways remain elusive and will be under a scope in the next research plan. We further raised a list of potential anti-HNSCC drug targets among genes in the TGF-β signaling pathway and/or the *HSF-1* activation network (Table S7) that may work in synergy with the neem extract against HNSCC. Although the potential effects of these drugs in synergy with the neem extract for HNSCC were not explored in this study, it is highly warranted to confirm their potential applicability in the clinical settings for HNSCC patients.

Despite the power of the pooled shRNA library screening, there have been caveats to the shRNA-based drug target discovery, and our study is no exception. Regarding the individual shRNA behavior in the shRNA screening, one would realize some biological discrepancies in comparison with the global gene expression and integrated pathway analyses. For example, although our pathway analyses identified the possible synergic interaction between neem extract and the TGF-β pathway, results from some of the individual hits in the shRNA screening were not concurrent. In the shRNA screening, knockdown of *SPHK1* and *LRRC15* (both are TGF-β pathway-related genes), for instance, were found to show no and positive effects on the cell growths of HSC-4 cells, respectively, under treatment of neem extract (Table S4). It can be hypothesized that the biological effects of the shRNA-mediated inhibition of single gene are heavily influenced by numerous experimental conditions: such as the time-scale (24hr in gene expression and six days in shRNA screening) and chemical activity (long-term culture in shRNA screening may inactivate the extract). One must also consider that single gene inhibition within a global web of signaling networks may result in various levels of different biological homeostasis in the cells, resulting in cellular phenotypes different from those expected from the function of the pathway itself. Also, any single gene has multiple of down-stream effectors, possibly influencing both activation and inhibition of cell growth simultaneously. Despite such limitations in our shRNA screening, we identified potential candidate pathways by utilizing an integrated gene set obtained from either or both of the gene expression and/or shRNA screening. This signifies the importance of combining multiple of global datasets and screening hits to identify candidate pathways of drug compounds with broad spectrums of biological effects such as the neem extract.

In summary, our study points out that neem leaf extract may offer a therapeutic potential to treat patients with HNSCC through its synergic modulations of pathways via *HSF-1* activation and TGF-β signaling. These finding will hopefully help developing novel anti-tumor drugs to be used with the neem leaf extract against HNSCC.

## Acknowledgements

The research is funded under the India-Japan Cooperative Science Programme by the Department of Science and Technology (DST), India to BP (Ref No. DST/INT/JSPS/P-181/2014), a grant from the Japanese Society of Promotion of Science (JSPS) to SI and funding from the Medical Research Institute, Tokyo Medical and Dental University for International Joint Research Project 2017. We thank PG Bharath Krishna for technical assistance.

## Author Contributions

VP, HK, MP and RS: data generation; NMK and HK: data analysis, NMK, HK, BP and SI: manuscript writing, BP and SI: Overall supervision and study guidance.

## Supporting Figure and Table legends

**Figure S1.**
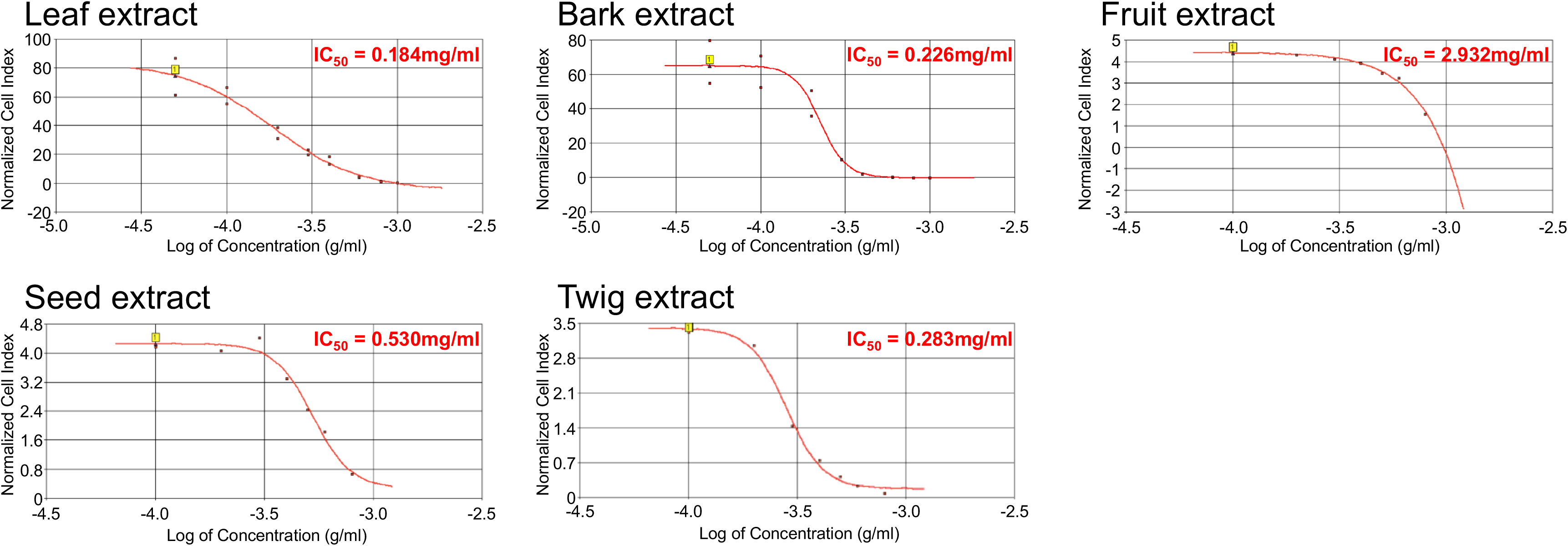
IC_50_ of neem extracts on HSC-4 cells.

**Figure S2.**
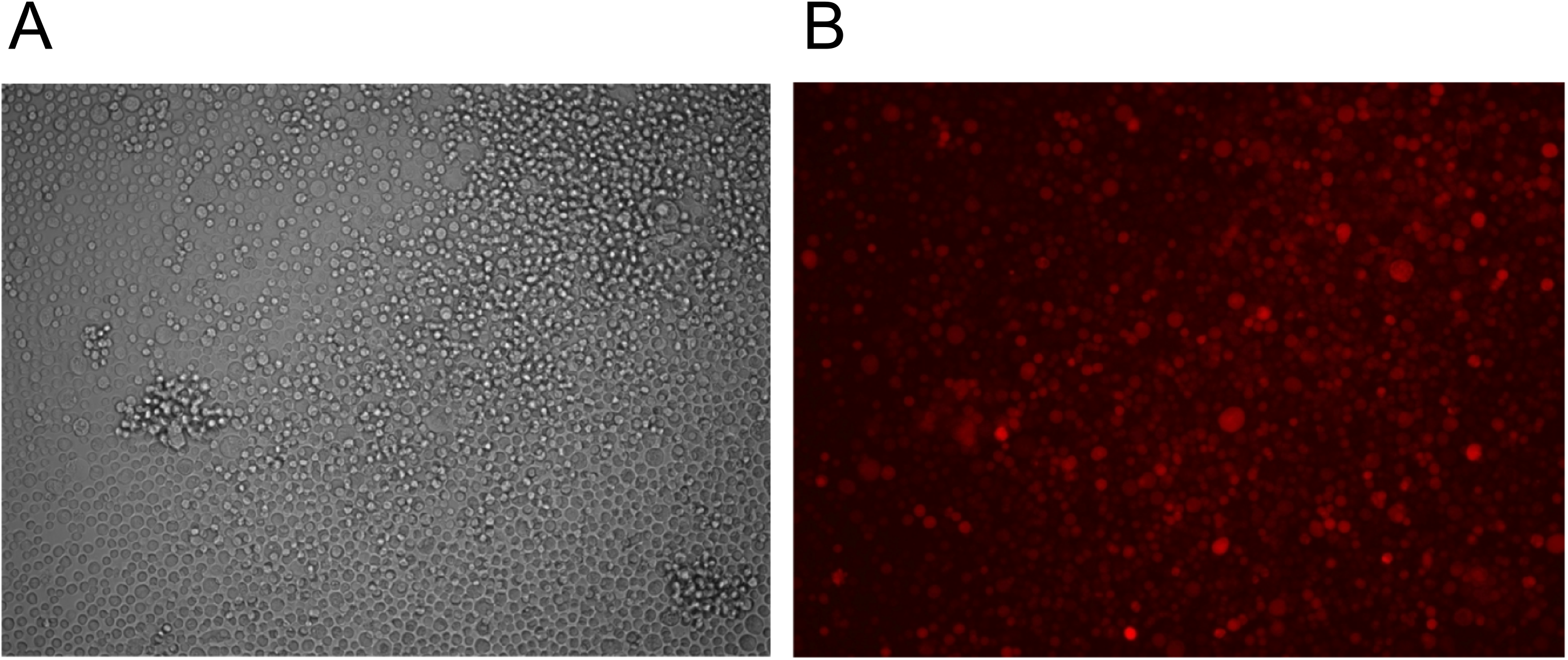
Cells after transduction (A) and the RFP signals at the time of harvesting (72 hr after the lentivirus infection) by using a fluorescent microscope (B).

**Table S1.** Expression values for all the genes at various time points and the follow-up rescue experiment along with the fold change statistics and cancer-specific pathways.

**Table S2.** Genes with altered expression across four or more time points.

**Table S3.** Results from the PCR amplification of the barcodes, in the control and in the leaf extract treated samples and the ratios representing clonal size changes.

**Table S4.** Changes in clonal size from shRNA experiments with leaf extracts for drug target analyses.

**Table S5.** Short-listed genes from the whole-genome gene expression (WGGX) assay and shRNA experiments for pathway analyses.

**Table S6.** Results from the pathway enrichment analyses performed using Reactome with a consensus list of 321 genes.

**Table S7.** Drug interactions with the genes perturbed in the HSF-1-based and the TGF-β signaling pathways.

